# *Prot-SpaM*: Fast alignment-free phylogeny reconstruction based on whole-proteome sequences

**DOI:** 10.1101/306142

**Authors:** Chris-Andre Leimeister, Jendrik Schellhorn, Svenja Schöbel, Michael Gerth, Christoph Bleidorn, Burkhard Morgenstern

## Abstract

Word-based or ‘alignment-free’ sequence comparison has become an active area of research in bioinformatics. While previous word-frequency approaches calculated rough measures of sequence similarity or dissimilarity, some new alignment-free methods are able to accurately estimate phylogenetic distances between genomic sequences. One of these approaches is *Filtered Spaced Word Matches*. Herein, we extend this approach to estimate evolutionary distances between complete or incomplete proteomes; our implementation of this approach is called *Prot-SpaM*. We compare the performance of *Prot-SpaM* to other alignment-free methods on simulated sequences and on various groups of eukaryotic and prokaryotic taxa. *Prot-SpaM* can be used to calculate high-quality phylogenetic trees from whole-proteome sequences in a matter of seconds or minutes and often outperforms other alignment-free approaches. The source code of our software is available through *Github*:

https://github.com/jschellh/ProtSpaM

## 1. Introduction

Evolutionary relationships between species are usually inferred by comparing homologous gene or protein sequences to each other. Here, groups of orthologous sequences have to be identified first, for which then multiple alignments are to be calculated. There are generally two different strategies of resolving phylogenies based on multiple alignments. In the so-called *supermatrix* approach, multiple sequence alignments of single genes or proteins are concatenated. A phylogenetic tree is inferred from the resulting matrix, *e.g.*, using *Maximum Likelihood* [56] or *Bayesian inference* [50]. Alternatively, gene or protein trees are inferred for every single multiple sequence alignment and the resulting phylogeny is inferred using *coalescent* models [40] or supertree [4] approaches.

All these steps are time consuming, and often manual intervention is required. Therefore, *word-based* or *alignment-free* alternatives have been proposed recently, which are much faster and which require much less data preparation. Most alignment-free methods compare the *word composition* of sequences [9, 16, 26, 52, 58, 62], with some approaches also considering background word frequencies [46, 47, 53, 63], see [48] for a review of these latter approaches. More recently, the *spaced-word* composition of sequences has been used for sequence comparison [27, 36, 42, 44]. Other alignment-free methods are based on the so-called *matching statistics*, that is they use the length of maximal common subwords [10, 61]. This has been extended to maximal common subwords with a certain number of mismatches [37, 45, 59, 60]. Alignment free approaches have been recently reviewed in detail [2, 22, 67].

Accurate alignment-free tools are urgently needed because of the huge volume of data generated by new sequencing techniques. Another advantage of alignment-free methods, compared to alignment-based approaches, is the fact that they can be applied to incomplete data, for example to unassembled sequencing reads or to partially sequenced genomes [14]. Note that some of the so-called ‘alignment-free’ approaches are based on comparing words of the input sequences to each other. So, strictly spoken, they are not ‘alignment-free’ since they align these words to each other. The term *alignment-free* is used nevertheless by most researchers, since these word-based approaches circumvent the need to calculate full pairwise or multiple alignments of the sequences under study.

The above mentioned approaches to alignment-free sequence comparison calculate ad-hoc measures of sequence similarity or dissimilarity. They are not based on stochastic models of molecular evolution, and they do not try to estimate distances between sequences in a statistically rigorous way. More recently, some alignment-free approaches have been proposed that are based on explicit models of DNA evolution. These methods are able to estimate the number of substitutions that have happened since two nucleic-acid sequences have evolved from their last common ancestor [11, 23, 24, 38, 41, 65].

A main application of alignment-free approaches is comparison of whole *genomes*. Consequently, most alignment-free methods have been designed to work on DNA sequences. If distantly related species are studied, though, phylogenetic trees are usually inferred from protein sequences rather than from DNA sequences. The reason for this is that protein sequences are more conserved than DNA sequences, as synonymous substitutions are not visible in proteins. Thus, for distal species, it may be hard to detect similarities between genes at the DNA-sequence level, while homologies may be still detectable among protein sequences. It is therefore highly desirable to have accurate alignment-free software tools that work on protein sequences, in addition to the available tools for DNA sequence comparison. Generic word-frequency methods can be applied to both DNA and protein sequences. As mentioned above, however, these methods do not estimate phylogenetic distances in a rigorous way. So far, there are no alignment-free approaches available that can accurately estimate evolutionary distances between protein sequences.

In this paper, we propose an alignment-free method which estimates the number of substitutions protein sequences since they evolved from their last common ancestor. Our approach is based on *Filtered Spaced Word Matches (FSWM)*, a concept we introduced recently for whole-genome sequence comparison [38], see [23, 65] for related approaches. We call the implementation of this new approach *<u>Prot</u>eome-based <u>Spa</u>ced-Word <u>M</u>atches (Prot-SpaM)*. The basic idea is to use gap-free pairwise alignments of fixed-length words with matching amino-acid residues at certain pre-defined positions. Such *spaced-word matches* can be rapidly identified and, after discarding random background matches, the remaining ‘homologous’ spaced-word matches can be used to estimate the phylogenetic distance between two taxa. To our knowledge, this is the first approach that accurately estimates evolutionary distances between protein sequences without the need to calculate full sequence alignments.

To evaluate our approach, we used simulated protein sequences and real-world whole proteomes. Test runs on the simulated sequences show that our distance estimates are very close to the true distances, for distance values of up to around 2.0 substitutions per sequence position. On the real-world sequences, we evaluated our approach indirectly, by phylogenetic analysis, as is common practice in the field. We used *Prot-SpaM* to estimate pairwise distances for various sets of taxa, and we applied the *Neighbor-Joining* algorithm [51] to calculate phylogenetic trees from the resulting distance matrices. These trees were finally evaluated by comparing them to reference trees that were determined by standard methods and can be considered to be reliable. We show that the trees obtained with our approach are generally very similar to the respective reference trees.

## 2. Method

*Prot-SpaM* is based on so-called *spaced-word matches* between sequences. For a binary pattern *P* of length *l* – *i.e.* for a word *P* over {0, 1} –, a *spaced-word match* with respect to *P* is a pair of words *w*_1_ and *w*_2_ of length *l* each, such that *w*_1_[*i*] = *w*_2_[*i*] for all *i ∈* {1, *…, l*} with *P* [*i*] = 1. Such positions *i* are called *match positions*, since the two words must have matching symbols at these positions. At positions *i* with *P* [*i*] = 0, the symbols *w*_1_[*i*] and *w*_2_[*i*] may differ; these positions are called *don’t-care positions*. A spaced-word match *between two sequences* is a spaced-word match involving one word from each of the two sequences. The number of *match* positions in a pattern or spaced-word match, respectively, is called its *weight w*. Below is an example for a spaced-word match between two toy protein sequences *S*_1_ and *S*_2_ with respect to a pattern *P* = 1100101 of length *l* = 7 and weight *w* = 4:

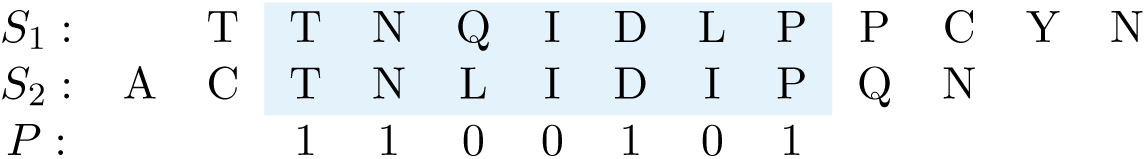

Similar to our original *FSWM* approach, we estimate distances between protein sequences based on selected spaced-word matches between them, with respect to one or several pre-defined patterns. Distance values are obtained by comparing the amino-acid residues that are aligned to each other at the *dont’-care* positions of the selected spaced-word matches. This is similar to estimating distances in standard alignment-based approaches – the only difference to those standard approaches is that we are using spaced-word matches instead of full sequence alignments.

To estimate distances in this way, one has to make sure that the matching spaced words that are selected, are *homologous* to each other, *i.e.* that they go back to the same origin in the last common ancestor of the two proteins that are compared. To distinguish such ‘homologous’ spaced-word matches from random background matches, we calculate a *score* for each spaced-word match using the BLOSUM62 substitution matrix [25]. Similar to the previous version of our program for nucleic-acid sequences, we define the *score* of a spaced-word match as the sum of substitution scores of the aligned amino acids at the don’t care positions of the underlying pattern *P*. Based on this score, our algorithm decides if a spaced-word match is homologous or not: if its score is below a certain threshold *T*, then a spaced-word match is considered a random match and is not further considered. As default we use a threshold value of *T* = 0. To see that this threshold accurately separates homologous from background spaced-word-matches, one can plot the *number* of spaced-word matches with a score *s* against *s*, see Figure 1; we call such a plot a *<u>Spa</u>ced-word-<u>M</u>atch hist<u>ogram</u>* or *SpaMogram*, for short. In these plots, two peaks are typically visible, a peak on the right-hand side representing *homologous* spaced-word matches and a peak on the left-hand side representing *background* matches; by default, our program uses patterns with a weight of *w* = 6 and with 40 *don’t-care* positions, *i.e.* with a length of *l* = 46.

**Figure 1:**
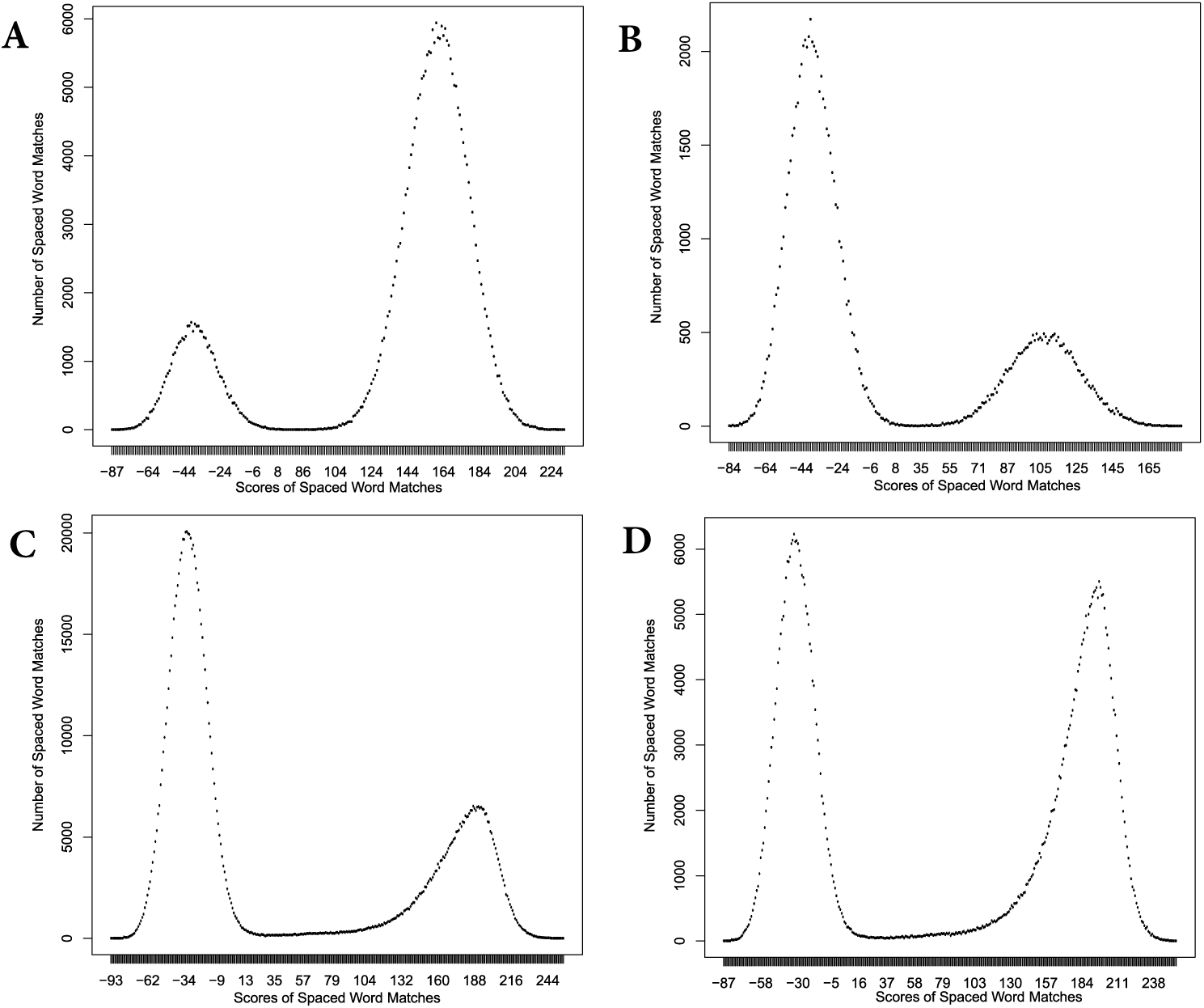
*Spaced-word histograms (SpaMograms)* for different data sets. (A) and (B) are based on simulated indel-free (concatenated) protein sequences with a total length of of 1.6 *×* 10^6^ amino-acid residues each, and with 0.3 (A) and 0.75 (B) substitutions per position, respectively. (C) and (D) are from a whole-proteome comparisons of plants, (C) comparing *Eucalyptus grandis* with *Capsella rubella* and (D) comparing *Gossypium raimondii* with *Carica papaya*.

Next, we make sure that we obtain a one-to-one mapping of spaced words in the two compared sequences. That is, we ensure, that each spaced word *w*_1_ in the first sequence, is matched to at most one spaced-word *w*_2_ in the second sequence, and vice versa, in our list of selected spaced-word matches. To achieve this, we use the same *greedy* algorithm that we described in our previous paper [38]. Finally, to estimate pairwise distances between the sequences we consider again the pairs of amino acids aligned to each other at the don’t care positions of the selected spaced-word matches and use the *Kimura* model to approximate the *PAM* distance between the sequences. To compare complete or partial proteomes to each other, our algorithm concatenates all protein sequences that belong to the same species. Special characters are inserted into the concatenated sequences to avoid spaced-word matches spanning more than one protein.

The accuracy and statistical stability of the described approach depends on the number of selected spaced-word matches: the more matches we obtain, the more accurate and stable the results of our method will be. To increase the number of spaced-word matches, the default version of our program uses *multiple* patterns, instead of one single pattern *P*. By default, our program uses a set of 5 patterns. To find good patterns sets, we integrated the tool *rasbhari* [20] into our implementation. *rasbhari* uses a *hill climbing* algorithm to reduce the *overlap complexity* [28] of pattern sets. Note that *rasbhari* uses a probabilistic algorithm to optimize pattern sets. It is therefore possible that, in different program runs, *rasbhari* returns different pattern sets, even if it is run with the same parameter values. Consequently, different runs of *Prot-SpaM* on the same sequences and with the same parameter setting can produce slightly different distance estimates.

## 3. Results

To assess the quality of our new approach and to compare it to other alignment-free methods, we used artificially generated as well as real-world protein sequences. For the test runs we used the default parameters of our program, namely 6 match positions and 40 don’t care positions – *i.e.* a total pattern length of 46 – and a threshold of *T* = 0 to discard background spaced-word matches. We compared our program to four other alignment-free methods that can be run on protein sequences, namely *ACS* [61], *FFP* [52], *kmacs* [37] and *CVTree* [46]. Since the original implementation of *ACS* is not publicly available, we used our own implementation of this approach by running *kmacs* with *k* = 0. The competing tools, too, were used with their default parameters. In addition to evaluating these tools on protein sequences, we ran *Filtered Spaced Word Matches* on the complete *genome* sequences of the same taxa.

### 3.1 Distance Estimation on Simulated Sequences

To evaluate the distances estimated by our program, we simulated sequences with the tool *pyvolve* [54]. *Pyvolve* simulates sequences along an evolutionary tree using continuous-time Markov models. It can use various substitution models such as *JTT* [29] and other models. Since there are no reliable stochastic models for insertions and deletions in protein sequences, the program produces indel-free sequences. We simulated pairs of sequences of length 100,000 with distances between 0 and 2 substitutions per position, in steps of 0.05, using the *JTT* model. To evaluate the estimated distance values, we generated 1,000 sequence pairs for each distance value and plotted the average of the estimated distances against the real *Kimura* distance of the respective sequence pairs, calculated with the program *protdist* from the *phylip* package [15]. To study the robustness of the estimated distances, we added error bars representing standard deviations to the plot. In addition to running *Prot-SpaM* with default parameters – *i.e.* with sets of five patterns –, we did a second series of test runs, where we used one single pattern. Figure 2 shows the results of these test runs.

**Figure 2:**
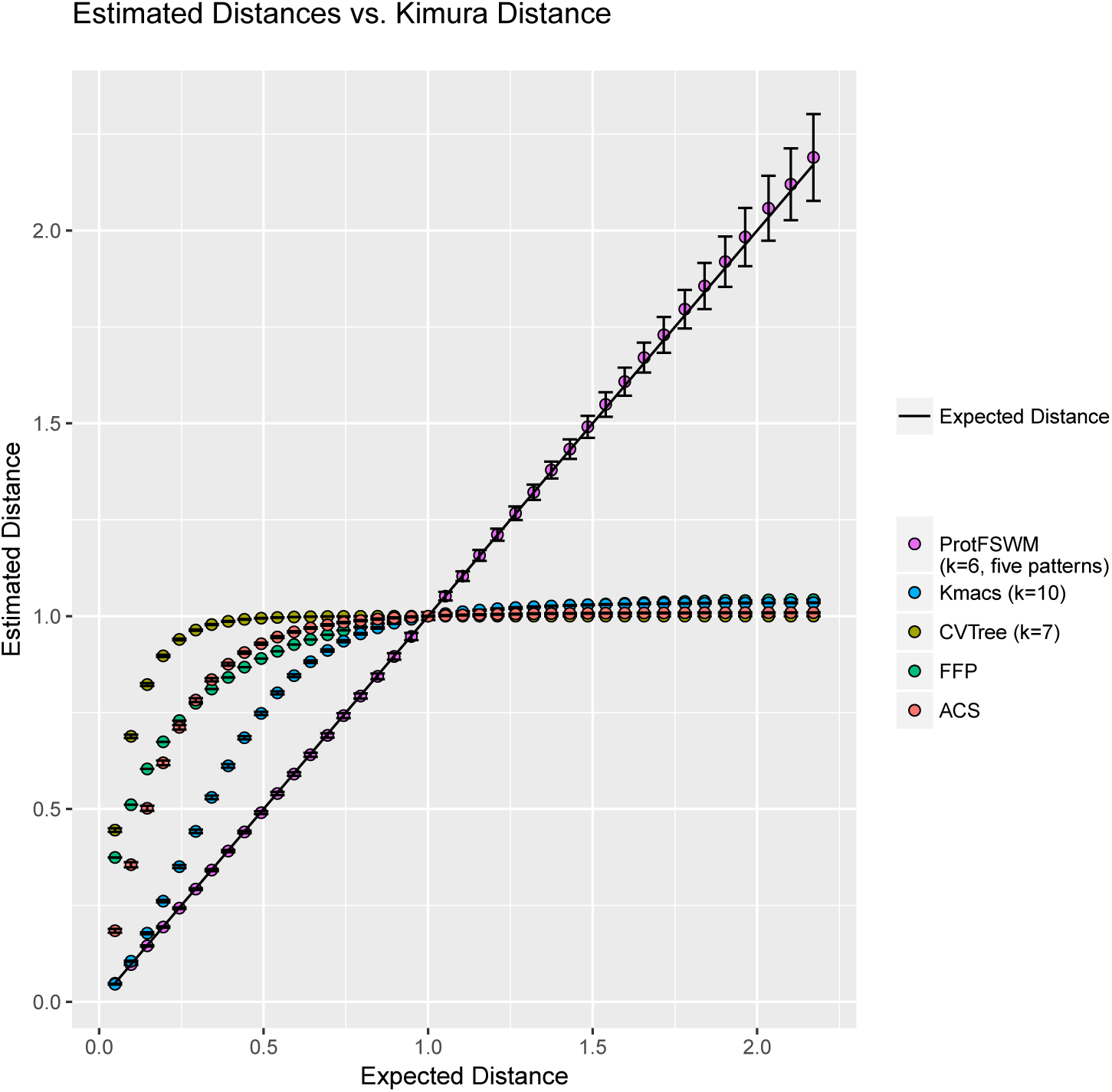
Distances calculated by *Prot-SpaM* and four other alignment-free methods calculated for pairs of simulated protein sequences, plotted against their distances calculated with the *Kimura* model. Error bars denote standard deviations. Note that *Prot-SpaM* estimates the number of substitutions per position since two sequences evolved from their lasts common ancestor – but *kmacs, CVTree, FFP* and *ACS* do not estimate distances in a rigorous way, but rather calculate some ad-hoc measure of sequence dissimilarity that is not a linear function of their real distances, *i.e.* the number of substitutions per position. Also, the absolute values of these distance measures are rather arbitrary for these four other programs. We therefore normalized the distances calculated by *kmacs, CVTree, FFP* and *ACS* such that they have a value of one for sequence pairs with a *Kimura* distance of one.

### 3.2 Phylogenetic tree reconstruction

Next, we applied the above five alignment-free methods to calculate phylogenetic trees from real-world protein sequences. For four different groups of species, we downloaded all available protein sequences from *GenBank* [1]; within each group, we calculated all pairwise distances between the species. We used the distance matrices obtained in this way as input for *Neighbor-Joining* [51] and compared the resulting trees to reference trees which we assume to reflect the respective correct phylogeny for each group. The *Robinson-Foulds* distances [49] between the reconstructed trees and the respective reference trees are shown in Table 1, the corresponding *branch score distances* [33] are in Table 2, and the program run times are in Table 3. Trees were visualized with *iTOL* [39]. *Neighbor-Joining* trees, *Robinson-Foulds* distances and *branch score* distances were calculated with the *phylip* package [15].

**Table 1:** *Robinson-Foulds* distances between trees generated with various alignment-free methods and the respective reference trees for various sets of taxa, see the main text for details. All programs were run on *protein* sequences or *whole proteomes*, respectively, except for *Filtered Spaced Word Matches (FSWM)* which was run on *whole-genome sequences* of the same species. Since the original implementation of *ACS* is not publicly available, we ran our own implementation, *kmacs*, with *k* = 0 instead.

| | *Prot-SpaM* | *FSWM* | *CVTree* | *FFP* | *kmacs*, $k = 10$ | *ACS* |
| --- | --- | --- | --- | --- | --- | --- |
| *E. coli* / *Shigella* | 4 | 6 | 24 | 40 | 42 | 38 |
| *Wolbachia I*, 252 proteins | 10 | 8 | 6 | 4 | 8 | 4 |
| *Wolbachia I*, whole proteomes | 6 | 6 | 8 | 16 | 8 | 12 |
| *Wolbachia II* | 28 | 20 | 44 | 54 | 26 | 16 |
| *Brassicea* | 0 | 0 | 6 | 8 | 2 | 6 |
| 813 prokaryotes | 1,020 | 1,348 | 886 | 1,452 | 880 | 960 |

**Table 2:**
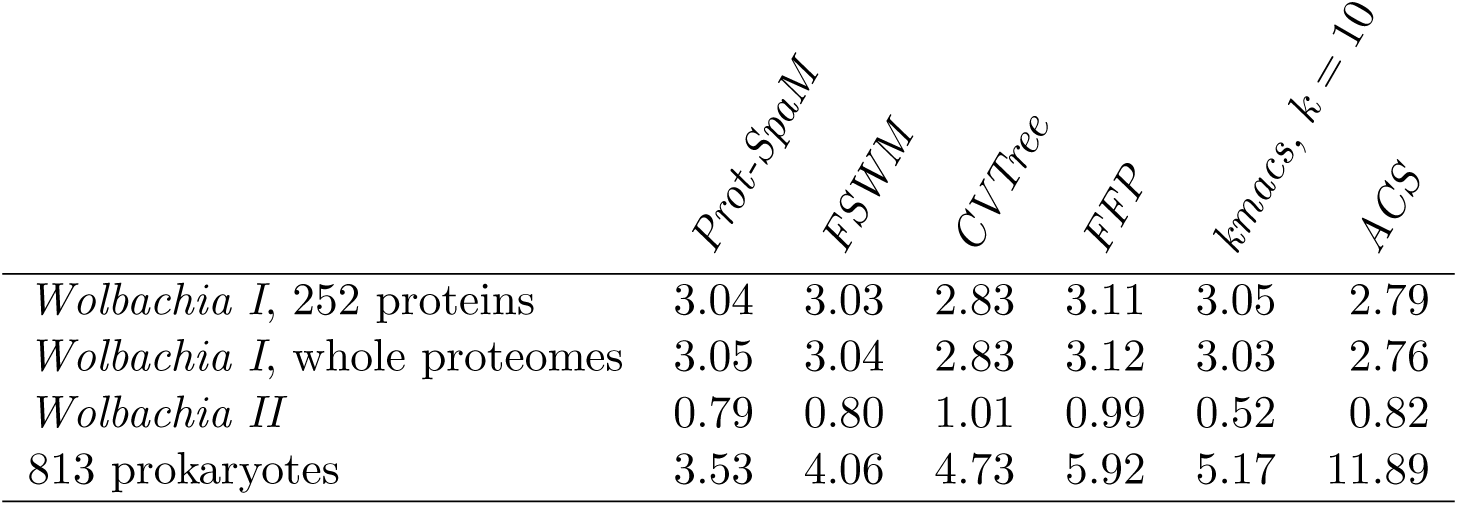
*Branch score* distances between trees generated with various alignment-free methods and the respective reference trees for various sets of taxa, see the main text and the legend of Table 1 for details.

**Table 3:**
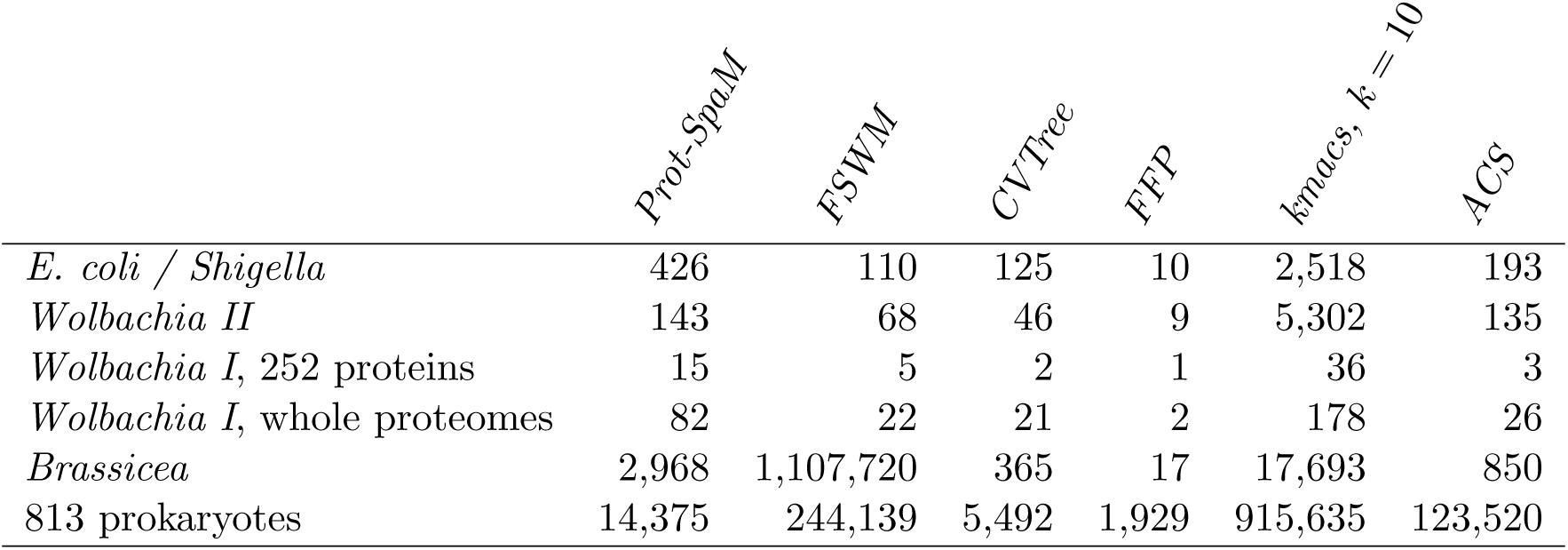
Program run time in seconds for different alignment-free approaches on our benchmark data sets.

#### E. coli / Shigella

Our first data set consists of 29 strains of *Escherichia coli* and *Shigella*. For each strain, we were able to download about 4,000-5,000 protein sequences; the total size of this data set is around 41 *mb*. Figure 3 shows the reference tree that we used and the tree obtained with the algorithm described in this paper. The reference tree was published by Zhou *et al.* [66] and is based on a multiple sequence alignment of 2,034 core genes and a *Maximum Likelihood* method. As can be seen in Table 1, our approach produced a tree with a topology almost identical to the reference tree; the *RF* distance between our tree and the reference tree was 4. The other alignment-free methods led to phylogenies with *RF* distances to the reference tree between 22 and 44.

**Figure 3:**
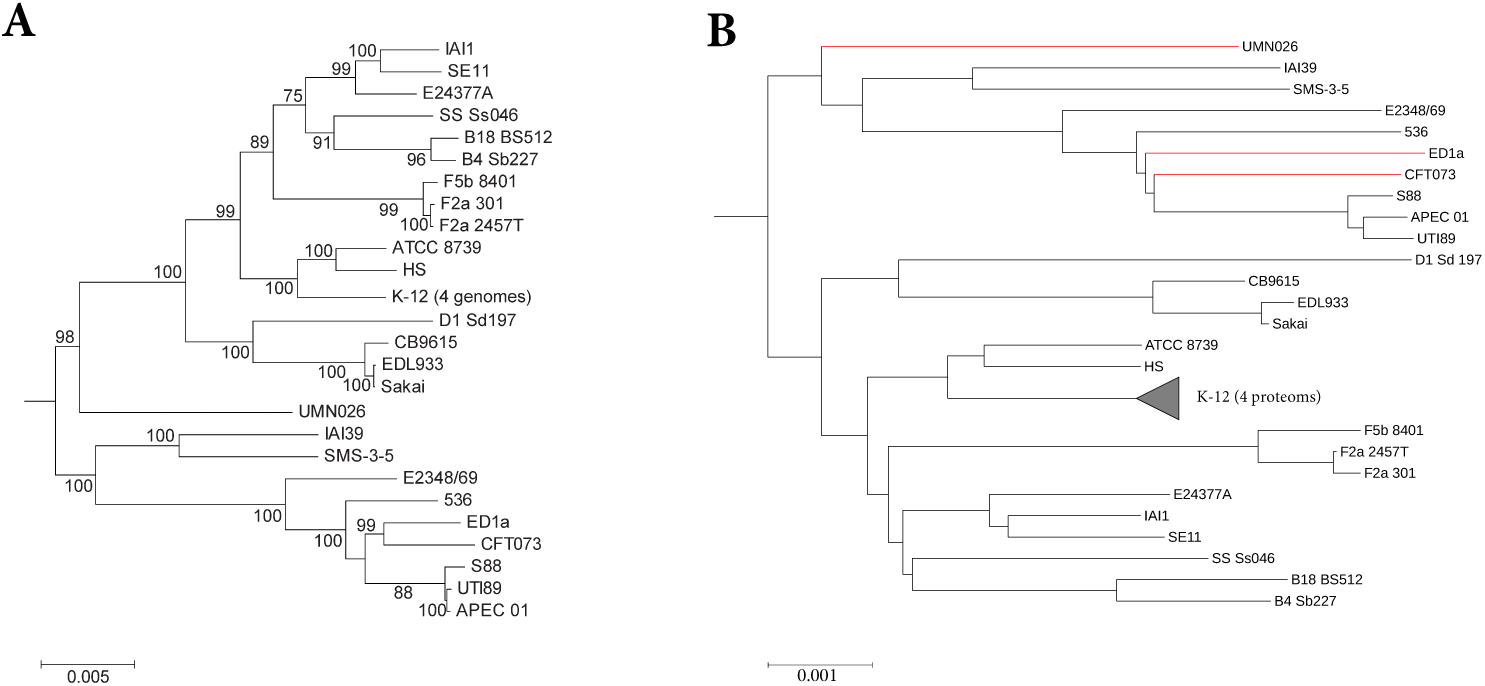
Reference tree (A) and tree calculated with *Prot-SpaM* with default parameters for a set of 29 *Escherichia coli* and *Shigella* strains. Differences in the topologies between the two trees are marked in red.

#### Wolbachia

As a second test case for benchmarking, we analysed the phylogeny of *Wolbachia* strains, a group of Alphaproteobacteria which are intracellular endosymbionts of arthropods and nematodes [64]. Within *Wolbachia*, 16 distinct genetic lineages (supergroups) are currently distinguished (named by capital letters A-F and H-Q), which may differ in host specificity and type of symbiosis [19]. We re-analyzed a phylogenomic dataset by [17], thereby focussing on relationships of strains within supergroups (*Wolbachia* I). For a second *Wolbachia* benchmarking dataset, we analysed relationships between supergroups based on available (draft) genomes, see below (*Wolbachia* II). For within supergroup relationships (*Wolbachia* I), a program run of *Prot-SpaM* on the whole proteome recovered a tree which is largely congruent in topology and branch lengths in comparison to a phylogenomic supermatrix analysis of 252 single-copy orthologs which excluded genes which showed signs of recombination. A comparison based on *RF* distances showed that our new method outcompetes other available alignment-free programs (Table 1). Interestingly, when only analysing the 252 ortholog dataset of [17] instead of whole proteomes, *RF* distances become bigger, and other alignment-free method perform better (Table 1). One interpretation is that misleading signal stemming from recombination events between *Wolbachia* strains is less problematic for alignment-free analysis then a reduction in he dataset size

Analysing relationships between supergroups has been historically regarded as difficult phylogenetic problem [5, 18]. Analysing all annotated proteins from available genomes with *Prot-SpaM* supported the monophyly of all supergroups. Moreover, this analysis found the same *Wolbachia* strains basally branching as recent analyses suggested. Surprisingly, the phylogenomic supermatrix analysis analysis of 252 single-copy orthologs which excluded genes which showed signs of recombination of this dataset recovered a topology which differs to previous study in not supporting the sistergroup relationship of supergroups A and B. In contrast, as found in previous analyses, the sistergroup relationship of supergroups A and B is supported by the *Prot-SpaM* analysis. The *Prot-SpaM* analysis also recovered some relationships between supergroups which differ from the topologies of our phylogenomic analysis or expectations from a recently published phylogenomic study [6]. However, it is known that supergroups differ in their base (and amino acid) composition, and it is currently unknown how this may impact alignment free methods. More sophisticated evolutionary models could alleviate these differences in future studies. Nevertheless, in this test case *Prot-SpaM* also outperforms other alignment free methods when comparing the resulting phylogenetic tree with a phylogenomic analyses based on a concatenated supermatrix (Table 1).

For the Wolbachia *II* dataset, we downloaded (if available) proteomes for all available *Wolbachia* draft and fully assembled genomes (47 in total) Proteins for *Wolbachia* strains which were lacking this information on *NCBI GenBank* were derived from translations using *GeneMark* version 2.5 [3]. We predicted groups of orthologous genes between these proteomes using *Orthofinder* version 2.1.2 [13] running under default parameters. Single copy genes present in all analysed strains (83 in total) were aligned using *MAFFT* version 7.271 with the L-INS-i algorithm [31], and tested for evidence of recombination using the pairwise homoplasy index (*PHI*) [7] with window sizes of 10, 20, 30, and 50. Recombining loci were subsequently removed from the dataset and the remaining loci concatenated using *FasConCat* version 1.0 [32]. The resulting supermatrix (68 loci, 20,787 amino acid positions) was subject to partitioned *Maximum Likelihood* analysis following best model and partition scheme selection in *IQ-TREE* version 1.6.2 [8, 30, 43];

#### Large-scale microbial phylogeny reconstruction

In 2013, J. Eisen’s group published a paper on the phylogeny of the microbial genomes that were available at the time [34]. As a basis of their study, they selected 24 single-copy marker genes and a non-redundant subset of taxa. To obtain such a subset, they used a greedy algorithm by M. Steel [57], making sure that marker genes from different taxa in the resulting subset had a distance to each other of at least 2 substitutions per 100 positions. This way, they obtained a non-redundant subset of 841 bacterial and archeal genomes from the more than 3,000 microbial genomes that were publicly available. Multiple sequence alignments of the marker genes were calculated with *hmmalign* [12] and were concatenated to a *supermatrix* which was used as input for the phylogeny programs *RAxML* [55] and *MrBayes* [50]. In addition, the authors used the Bayesian tree-reconciliation program *BUCKy* [35] to the same set of marker genes. The trees they obtained with these different methods were found to be similar to trees obtained based on *16S RNA* genes.

To evaluate *Prot-SpaM*, we used the 841 microbial genomes from Lang *et al.* and downloaded all protein sequences from these taxa that were available through *GenBank*. For 28 out of the 841 taxa, we were unable to obtain protein sequences, so we obtained a slightly reduced subset of 813 taxa, compared to the taxa used by Lang *et al*. First, we applied *Prot-SpaM* to all available protein sequences from these 813 taxa. In addition, we ran *Prot-SpaM* on the protein sequences encoded by the 24 marker genes from Lang *et al.* and, finally, we applied our previous approach *Filtered Spaced Word Matches* [38] to the 841 genome sequences. The trees that we obtained with our different alignment-free approaches are shown in Figure 4, together with the *Maximum Likelihood* tree from [34] which we considered as a reliable reference. Clades from this reference tree are color-coded in Figure 4. As can be see from the color coding, the tree obtained with *Prot-SpaM* from the available protein sequences contains essentially the same clades as the reference tree. There are some differences within the clades, though, that should be further investigated (J. Eisen, personal communication).

**Figure 4:**
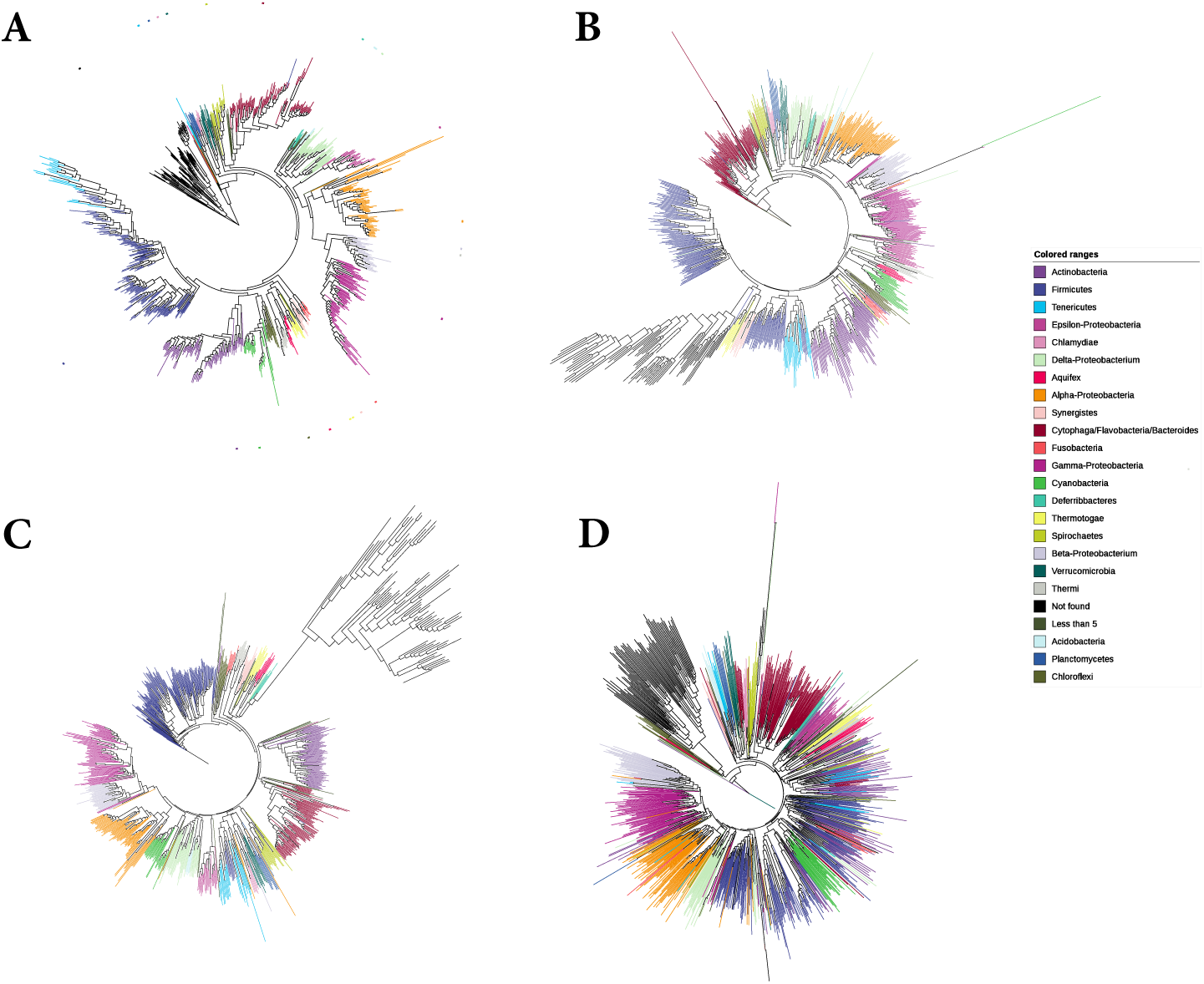
Phylogenetic trees for a set of 841 microbial taxa studied by Lang *et al.* [34]. (A) Maximum-Likelihood tree constructed by Lang *et al.* based on a super alignment of 24 selected genes, (B) tree constructed with our approach, as described in this paper, for 813 taxa for which the proteomes are available in *GenBank*, (C) tree constructed with our approach based on the proteins corresponding to the 24 genes selected by Lang *et al.* and (D) tree reconstructed using our program *FSWM* [38] on the 841 whole-genome sequences.

#### Brassicales

Finally, we used a set of plant taxa [21] that we had already used in previous studies to evaluate alignment-free approaches to genome sequence comparison [36–38]. The data set that we used in these previous papers consisted of 14 brassicales species. In *GenBank*, however, the proteomes could be downloaded only for 11 of the 14 species, so we had to limit our test runs to these 11 species. Figure 5 shows the reference tree of the 14 original species, together with trees of the 11 species with available proteomes, calculated with the alignment-free methods that we evaluated in this paper.

**Figure 5:**
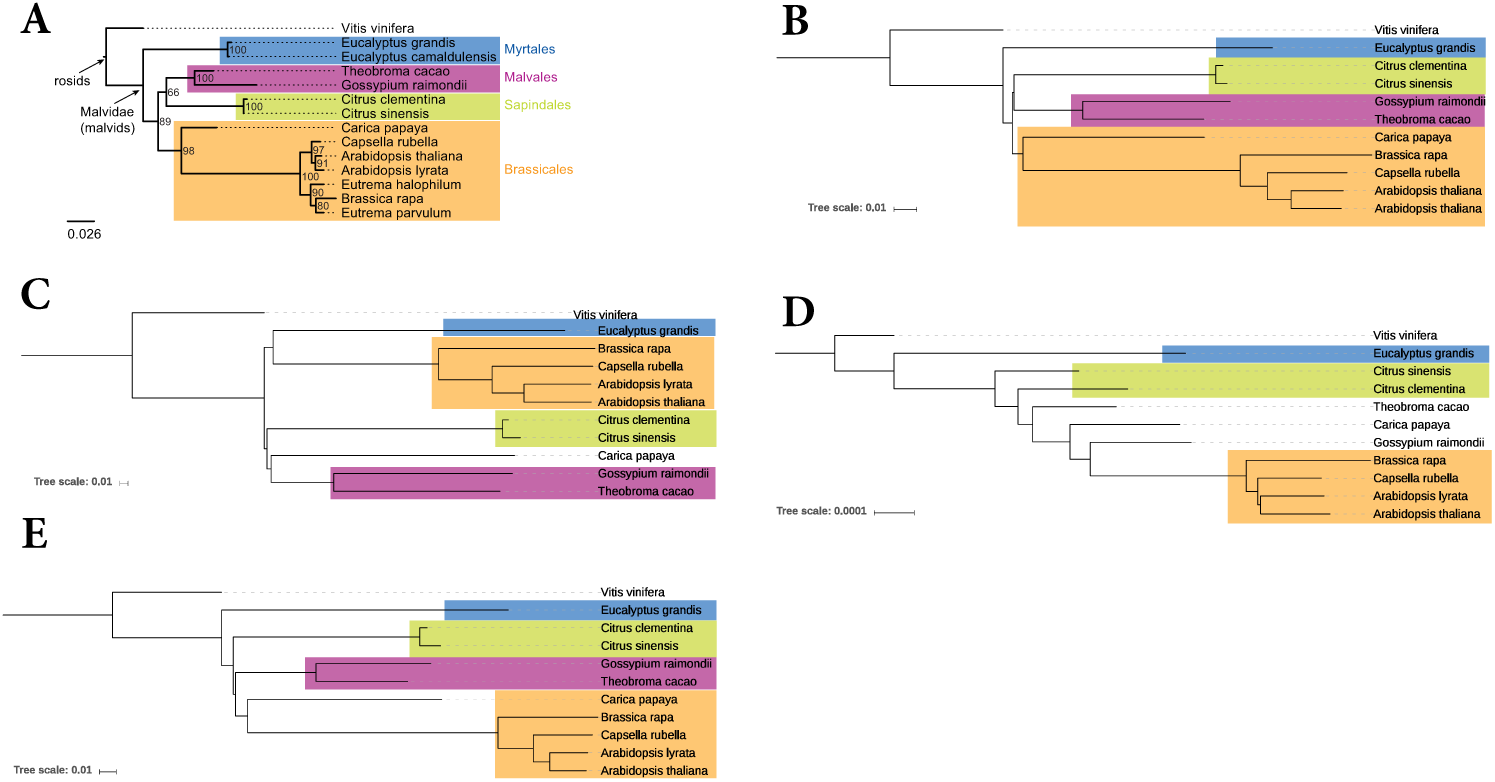
Phylogenetic trees of plant taxa. (A) reference tree from [21], and trees constructed with (B) the approach described in this paper, (C) *ACS* [61], (D) *FFP* [52], and (E) *kmacs* [37]. The original data set contained 14 taxa, but only for 11 taxa, the proteomes could be downloaded through *GenBank*. For completeness, we show the reference for all 14 taxa.

## 4. Discussion

A number of so-called ‘alignment-free’ approaches have been proposed in recent years to estimate phylogenetic distances between genome sequences, *i.e.* to estimate the average number of substitutions per position that have occurred since two genomes have evolved from their last common ancestor.

One of these approaches is *Filtered-Spaced Word Matches (FSWM)* [38]. Some of the existing alignment-free approaches can also be applied protein sequences. A draw-back of these methods is that they calculate only rough measures of sequence similarity or dissimilarity, they do not estimate distances in terms of events that may have occurred during evolution. In this study, we introduced *Prot-SpaM*, a new implementation of *FSWM* that can compare complete or incomplete *proteomes* to each other. To our knowledge, *Prot-SpaM* is the first tool that can accurately estimate phylogenetic distances between protein sequences without the need to calculate full sequence alignments.

Our benchmark results show that distance estimates obtained with our approach are accurate for a large range of phylogenetic distances. Distances between protein sequences calculated with *CVTree, ACS, FFP* and *kmacs*, by comparison, are monotonously increasing with the number of substitutions between the compared sequences. These curves, however, are far from linear, and they flatten out at distance values somewhere between 0.5 and substitutions per position, see Figure 2. By contrast, *Prot-SpaM* estimates distances with high accuracy for up to around 2.0 substitutions per position. For higher distance values, the calculated distances become less stable, as can be seen from the error bars in Figure 2. Moreover, for large distances, our program tends to slightly overestimate distances.

Phylogenetic trees generated from these distance values are generally of high quality. Table 1 shows that, for various sets of taxa, trees based on *Prot-SpaM* distances, calculated from whole proteomes, are more similar to the respective reference trees than the trees that we obtained with alternative alignment-free methods. Interestingly, this result was reversed for the taxa set *Wolbachia I*, when we used only proteins from 252 selected genes instead of all available protein sequences. Another interesting result is the performance of *Prot-SpaM*, compared to our corresponding previous approach that works on genome sequences. For most groups of taxa in our study, the results of *Prot-SpaM* and *FSWM* were of similar quality, in the sense that the *RF* distances to the reference trees were comparable for both approaches. However, for the set of 813 prokaryote taxa, our new spaced-words approach performed better on whole-proteomes than our previous approach on whole genomes, as is shown in Figure 4 and Table 1. This discrepancy is most likely due to the large phylogenetic distances in this data set; at these distances, homologies are generally better detectable at the protein level than at the *DNA* level.

The main advantage of our new approach is its high speed, with only a small loss of quality, compared to more traditional, alignment-based approaches to phylogeny reconstruction. A program run of our software on the set *Wolbachia II* that consists of the proteomes of 47 taxa, took around three minutes. Moreover, our approach can reliably distinguish between local homologies and random background spaced-word matches. Therefore, it can be applied to complete or incomplete proteomes, it is not necessary to select orthologous genes or proteins in a first step. For the taxa set *Wolbachia I*, we obtained better results when we used all available protein sequences from GenBank, than with the proteins corresponding to 252 carefully selected genes. Therefore, we think that *Prot-SpaM* should be a useful addition to existing approaches to phylogeny reconstruction.

## Acknowledgements

We thank Jonathan Eisen for his comments on the phylogeny of the 841 microbial taxa that were studied in Lang *et al.* We acknowledge support by the *Open Access Publication Funds* of the Göttingen University.

